# Optical mapping compendium of structural variants across global cattle breeds

**DOI:** 10.1101/2022.05.05.490773

**Authors:** A. Talenti, J. Powell, D. Wragg, M. Chepkwony, A. Fisch, B.R. Ferreira, M.E.Z. Marcadante, I.M. Santos, C.K. Ezeasor, E.T. Obishakin, D. Muhanguzi, W. Amanyire, I. Silwamba, J.B. Muma, G. Mainda, R.F. Kelly, P. Toye, T. Connelley, J. Prendergast

**Author notes:** Contributed equally (T. Connelley, J. Prendergast).

## Abstract

Structural variants (SV) have been linked to important bovine disease phenotypes, but due to the difficulty of their accurate detection with standard sequencing approaches, their role in shaping important traits across cattle breeds is largely unexplored. Optical mapping is an alternative approach for mapping SVs that has been shown to have higher sensitivity than DNA sequencing approaches. The aim of this project was to use optical mapping to develop a high-quality database of structural variation across cattle breeds from different geographical regions, to enable further study of SVs in cattle.

To do this we generated 100X Bionano optical mapping data for 18 cattle of nine different ancestries, three continents and both cattle sub-species. In total we identified 13,457 SVs, of which 1,200 putatively overlap coding regions. This resource provides a high-quality set of optical mapping-based SV calls that can be used across studies, from validating DNA sequencing-based SV calls to prioritising candidate functional variants in genetic association studies and expanding our understanding of the role of SVs in cattle evolution.

## Background & Summary

Structural variants (SV) are a heterogeneous class of genetic variants involving large fragments of the genome (>50bp)^1^. These variants include genomic insertions and deletions (InDels), inversions, duplications, translocations and more complex rearrangements^2^. Single nucleotide polymorphisms (SNPs) have been the primary focus of studies trying to map genetic loci underlying important cattle phenotypes. However, there are multiple lines of evidence suggesting SVs likely underlie many important cattle traits. As many as 25-29% of all protein truncating events are thought to be caused by an SV in humans^1^ and notably, despite being less well studied, SVs have already been tied to key livestock phenotypes. For example, a duplication of the CIITA class II major histocompatibility complex transactivator gene in cattle has been tied to resistance to intestinal nematodes^3^ and a 12Kb copy number variant has been linked to mastitis in cattle^4^. Chromosomal translocations and duplications have been linked to skin pigmentation, a phenotype closely tied to environmental adaptation, and SVs across livestock species have been linked to phenotypes such as olfaction or resistance to adenocarcinoma-causing viruses^5^. Importantly SVs are responsible for approximately 5-10 times as many heritable nucleotide sequence differences between individuals than SNPs^6^. Unlike SNPs, that only effect a single basepair, and most often far from coding regions, SVs effect large regions and potentially multiple genes. Consequently, although smaller in number, any given novel SV event is more likely to have a phenotypic consequence.

The two most popular methods used to detect SVs are high-throughput sequencing (HTS) and array comparative genomic hybridisation (aCGH), both of which have been applied to European cattle^7–9^, but with few studies performed in other cattle breeds^10–12^. Each technology has advantages and limitations. aCGH, for example, involves measuring binding to probes covering the reference genome, and therefore it can only detect relative copy number changes between sample pairs and cannot for example detect novel insertions. Resolution is also limited. A major advantage of HTS approaches is that theoretically they can detect SVs at base-pair resolution. However, accurate calling of SVs from HTS data has proven to be difficult for a number of reasons including poor reference assemblies, chimeric reads, aligners penalising reads that don’t match the reference and the difficulties of sequencing and mapping to repetitive regions. This is exemplified by the generally poor agreement between SV callers even when run across the same samples^13,14^. Approaches using long reads and *de novo* assembly can still have true positive rates as low as 77%, even when using simulated data^15^.

Optical mapping (OM), a light microscope-based method that labels and physically locates specific motifs in the genome^16^, offers an alternative protocol to accurately detect large SVs. OM molecules can be consistently hundreds of Kb long, allowing for the detection of complex rearrangements undetectable using HTS. Despite the limitation of not being able to detect the actual sequence of the identified SVs, as well as missing smaller SVs, OM has a very high sensitivity and specificity, allowing for the generation of high-quality catalogues of SVs in individuals^17^. A study in humans successfully used OM reads to identify SVs in a total of 26 genomes revealing population-specific patterns of structural variation^18^.

In this study, we generated the first catalogue of cattle OM data for 18 animals from 9 different global breeds, and three continents, to better characterise common SVs across the cattle pan-genome. This data is a particularly valuable resource of SVs for the cattle species to intersect with other datasets, for example, for the validation of SV calls from other approaches.

## Methods

### Sample preparation

We selected a set of 18 cattle across 9 divergent European, African and Indian breeds representative of Indicine, Sanga and Taurine ancestries (Table 1). Blood was collected by jugular venipuncture into EDTA vacutainers. Somatic recombination in B cells and T cells means the Ig and TCR loci in these cell types will be highly heterogenous, confounding accurate reconstruction of germline SVs at these loci from whole blood samples. Consequently, after the erthyrocyte lysis, monocytes were purified from the leukocytes using a MACS positive selection protocol with an anti-bovine SIRPα mono-clonal antibody (ILA-24^19^). Agarose plugs containing 5 x10^5^ – 1×10^6^ of isolated monocytes were prepared using the Bionano Blood and cell culture DNA isolation kit (Bionano Genomics, San Diego, US) according to the manufacturer’s instructions and the extracted DNA used for analysis on the Bionano Saphyr platform to generate ~100X optical mapping coverage of each genome.

**Table 1:** Description of the samples. Table describing the breeds and ancestry of samples, with the continent and country of origin. The identifiers, as well as the ENA accession codes, for each of the two animals sampled per breed are also reported.

| Sampling<br>Continent | Sampling<br>Country | Group | Breed | ENA project<br>ID | ENA sample<br>ID |
| --- | --- | --- | --- | --- | --- |
| S.<br>America | Brazil | Indicine | Nelore | PRJEB52551 | ERS11891755 |
|  |  |  |  | PRJEB52551 | ERS11891754 |
| Africa | Kenya | Indicine | Boran | PRJEB52551 | ERS11891767 |
|  |  |  |  | PRJEB52551 | ERS11891766 |
| Africa | Nigeria | Indicine | White<br>Fulani | PRJEB52551 | ERS11891768 |
|  |  |  |  | PRJEB52551 | ERS11891769 |
| Africa | Zambia | Indicine | Angoni | PRJEB52551 | ERS11891764 |
|  |  |  |  | PRJEB52551 | ERS11891765 |
| Africa | Uganda | Sanga | Ankole | PRJEB52551 | ERS11891756 |
|  |  |  |  | PRJEB52551 | ERS11891757 |
| Africa | Zambia | Taurine | Barotse | PRJEB52551 | ERS11891762 |
|  |  |  |  | PRJEB52551 | ERS11891763 |
| Africa | Nigeria | Taurine | N'Dama | PRJEB47998 | ERS8452869 |
|  |  |  |  | PRJEB47998 | ERS8452868 |
| Europe | United Kingdom | Taurine | Hereford | PRJEB52551 | ERS11891760 |
|  |  |  |  | PRJEB52551 | ERS11891761 |
| Europe | United Kingdom | Taurine | Holstein-Friesian | PRJEB52551 | ERS11891759 |
|  |  |  |  | PRJEB52551 | ERS11891758 |

### Bionano Solve optical mapping processing

OM reads were filtered using the filter_SNR_dynamic.pl script with default parameters included with the Solve workflow, and then processed through the Bionano Solve^20^ pipeline (v3.3_10252018) using two different releases of RefAligner to overcome bugs preventing the successful assembly of the reads (version 7915.7989rel and 10330.10436rel). We generated the reference CMAP for the ARS-UCD1.2 genome with the Y chromosome from the 1000 bulls genome project (https://sites.ualberta.ca/~stothard/1000_bull_genomes/) using fa2cmap_multi_color.pl (default options and specifying the DLE1 as enzyme). The resulting data were imported into the Bionano Access (v1.6) software, and single-sample SVs were filtered using the recommended thresholds for SVs generated using Bionano Solve prior to v1.6.0 with the sizes recommended to achieve 90% sensitivity (see https://bionanogenomics.com/wp-content/uploads/2018/04/30110-Bionano-Solve-Theory-of-Operation-Structural-Variant-Calling.pdf): minimum insertion size of 5Kb, minimum deletion size of 5Kb, minimum inversion size of 100Kb, and minimum duplication size of 150kb.

Filtered smap format files were converted to vcf format using smap_to_vcf_v2.py and sorted with bcftools (v1.10.2^21^). The resulting SVs were screened using bcftools and retained if 1) they had successfully been genotyped, 2) their size was >1Kb and 3) their quality was >= 20. The latter filtering largely removed all translocations, duplications, and complex events due to these having either very low (<1) or nil quality values.

We then defined the total amount of non-redundant reference sequence involved in a high-quality deletion. For each deletion, we calculated the central point in the genomic region affected by the SV:

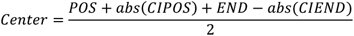

Where POS is the initial position, END is the end position, CIPOS is the confidence interval of POS and CIEND is the confidence intervals of END. Having defined the central point of the region, we defined the initial and final positions of the SV as:

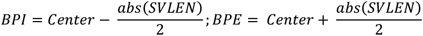

Where BPI and BPE are the limits of the SV and SVLEN is the size of the SV.

We then concatenated the regions for all the individuals, sorted them and merged them using bedtools sort and bedtools merge^22^ to remove any redundancies among the regions. Following filtering, we merged the resulting variants within samples using SURVIVOR (v1.0.7^2^) accounting for the SV type and collapsing those whose break points were within 1kb. We represented the intersection of SVs across individuals by extracting the support vectors generated by SURVIVOR^2^ at merging time, and plotted them using the UpSet function from the R^23^ package ComplexHeatmap^24^ (v2.8.0). We extracted the support value (i.e. how many animals present a specific SV) and SV size for each variant in the combined VCF and tested whether the SVs found in one individual only (support = 1) were significantly larger than those shared among individuals (support > 1) by performing a Wilcoxon signed-rank test in a custom R script.

Finally, we defined which of the final set of SVs were found to potentially affect a gene. We ran VEP v105^25^ to predict which SVs were likely to disrupt a gene’s function, with the options --sift b (both preditions score and term), --nearest symbol (report the gene symbol), and --distance 200 (200 bp up and downstream consequence prediction). Those variants presenting coordinates referring to the negative strand (end position smaller than initial position) were manually fixed through an in-house script. We then investigated which SVs putatively overlap a coding region annotated in the cow genome by intersecting merged SVs with coding sequence intervals. Intersecting genes were investigated with FUMA^26^ to identify enriched gene ontologies and gene sets using all 35,142 gene elements with a unique Entrez gene ID as the background list.

## Data Records

The datasets presented here are stored at ENA under analysis IDs PRJEB47998 and PRJEB52551. The data are uploaded in Bionano BNX format compatible with downstream analyses. The output of the Solve workflows can be downloaded from Zenodo (DOIs: 10.5281/zenodo.6516993 and 10.5281/zenodo.6517172). The raw VCF files, converted using smap_2_vcf_v2.py, can be found on Zenodo with DOI: 10.5281/zenodo.6516933.

## Technical Validation

### Assembly statistics and SV calling

We aligned the Saphyr optical mapping reads to the ARS-UCD1.2 genome^27^, expanded with the BTau5 Y chromosome generated by the 1000 bulls genome project, using Bionano Solve (v3.3 and 3.5) to assemble the genome maps and call SVs. The two NDama samples had previously been used to validate SV using graph genome approaches^28^.

Workflow metrics are provided in Supplementary Data 1, summarising key metrics for the analysis of these samples in comparison to the recommended values from Bionano (see https://bionanogenomics.com/wp-content/uploads/2017/03/30255-Bionano-Access-Assembly-Report-Guidelines.pdf).

Unfiltered molecules had average read lengths of 131.9-219.8kb (recommended >150kb for both metrics) and molecule N50s ranged from 185.2-361.9kb across the samples (recommended >150Kbp). Following molecule filtering, all samples were within the recommended average length (245.5-383.1, recommended >230Kbp) and molecule N50 (245.0-426.5, recommended >230Kbp), and only 1 sample (Angoni 1) was slightly below the recommended label density (13.1-16.4, recommended 14-17). Importantly all samples passed the recommended values for the effective coverage of the reference (72.5-128.2, recommended >70) and of average confidence (30.1-43.2, recommended >20).

All samples also generated assemblies with high genome map N50s for both the diploid (71.7-85.0, recommended >50) and haploid (71.3-84.5, recommended >50) assemblies. Despite the low proportion of assembled contigs aligning to the reference genome (0.14-0.25, recommended >0.70), the high uniquely aligned length by reference length (0.835-0.906, recommended >0.85) shows the presence of long assembled contigs. The contigs present a high fraction of molecules aligned (0.77-0.94, recommended >0.80), effective coverage assembly (37.7-66.7, recommended >40) and average confidence (38.5-51.3, recommended >20).

Overall, 1 sample had 11 metrics within the recommended values, 6 had 12 metrics within the recommended values, 9 had 13 metrics within the recommended values and 2 had 14 metrics within the recommended values.

The Bionano Solve workflow identified a number of SV in each sample, ranging from 4,944 to 11,184 for a Hereford and Nelore, respectively (Table 2). This mirrors the evolutionary distance of each sample from the reference genome, with the European taurine possessing fewer SVs (4,944-5,652) than the other samples, and an African taurine N’Dama possessing the least among the non-European individuals (N = 6,254). Relative SV numbers consequently broadly mirror prior expectations.

**Table 2:** Raw number of structural variants (SVs) and type detected in the different samples. This table summarises the number of raw SVs detected in each sample, and their classification (e.g. insertion, deletion, duplication, inversion and inter- and intra-chromosomal translocation).

| Sample | Dels | Insert. | Dups. | Inversion breakpoints | Interchr. translocation breakpoints | Intrachr. translocation breakpoints | Total |
| --- | --- | --- | --- | --- | --- | --- | --- |
| Angoni 1 | 4349 | 4505 | 45 | 91 | 13 | 4 | 9007 |
| Angoni 2 | 4387 | 4673 | 64 | 100 | 11 | 7 | 9242 |
| Ankole 1 | 4314 | 4324 | 67 | 111 | 10 | 8 | 8834 |
| Ankole 2 | 3911 | 3984 | 66 | 101 | 8 | 5 | 8075 |
| Barotse 1 | 3971 | 4044 | 42 | 52 | 11 | 6 | 8126 |
| Barotse 2 | 4199 | 4159 | 67 | 106 | 7 | 9 | 8547 |
| Boran 1 | 4935 | 5087 | 56 | 113 | 15 | 6 | 10212 |
| Boran 2 | 4990 | 5007 | 68 | 138 | 6 | 14 | 10223 |
| Hereford 1 | 2465 | 2380 | 43 | 41 | 11 | 4 | 4944 |
| Hereford 2 | 2435 | 2437 | 77 | 88 | 9 | 7 | 5053 |
| Holstein 1 | 2756 | 2759 | 48 | 52 | 15 | 18 | 5648 |
| Holstein 2 | 2702 | 2801 | 59 | 76 | 9 | 5 | 5652 |
| N'Dama 1 | 3411 | 3481 | 92 | 125 | 10 | 13 | 7132 |
| N'Dama 2 | 3005 | 3082 | 67 | 86 | 6 | 8 | 6254 |
| Nelore 2 | 5294 | 5508 | 58 | 113 | 11 | 5 | 10989 |
| Nelore 1 | 5420 | 5499 | 96 | 136 | 15 | 18 | 11184 |
| White Fulani 1 | 4467 | 4642 | 54 | 114 | 11 | 3 | 9291 |
| White<br>Fulani 2 | 4782 | 4805 | 41 | 45 | 17 | 14 | 9704 |

### Variant statistics

SVs were filtered using Bionano Access, excluding SVs with unknown dosages, and retaining those larger than 1Kb and with a quality >20. SVs for each individual were then combined using SURVIVOR^2^ if the breakpoints were within 1Kb, i.e. below the effective resolution of the approach^2^. This process allowed us to select a catalogue of 13,457 SVs across the genome, containing 8,262 insertions, 5,191 deletions and 4 inversions (see Supplementary Data 2 and 3 for the details on the type of SV identified). No duplications, inverted duplications and translocations passed the quality filtering. Consistent with results from previous studies^28^, most of the post-filtering insertions and deletions identified fell into the smaller classes, though 1,796 SVs (403 deletions, 1,389 insertions and 4 inversions) of over 50kb in length were identified (Figure 1).

**Figure 1:**
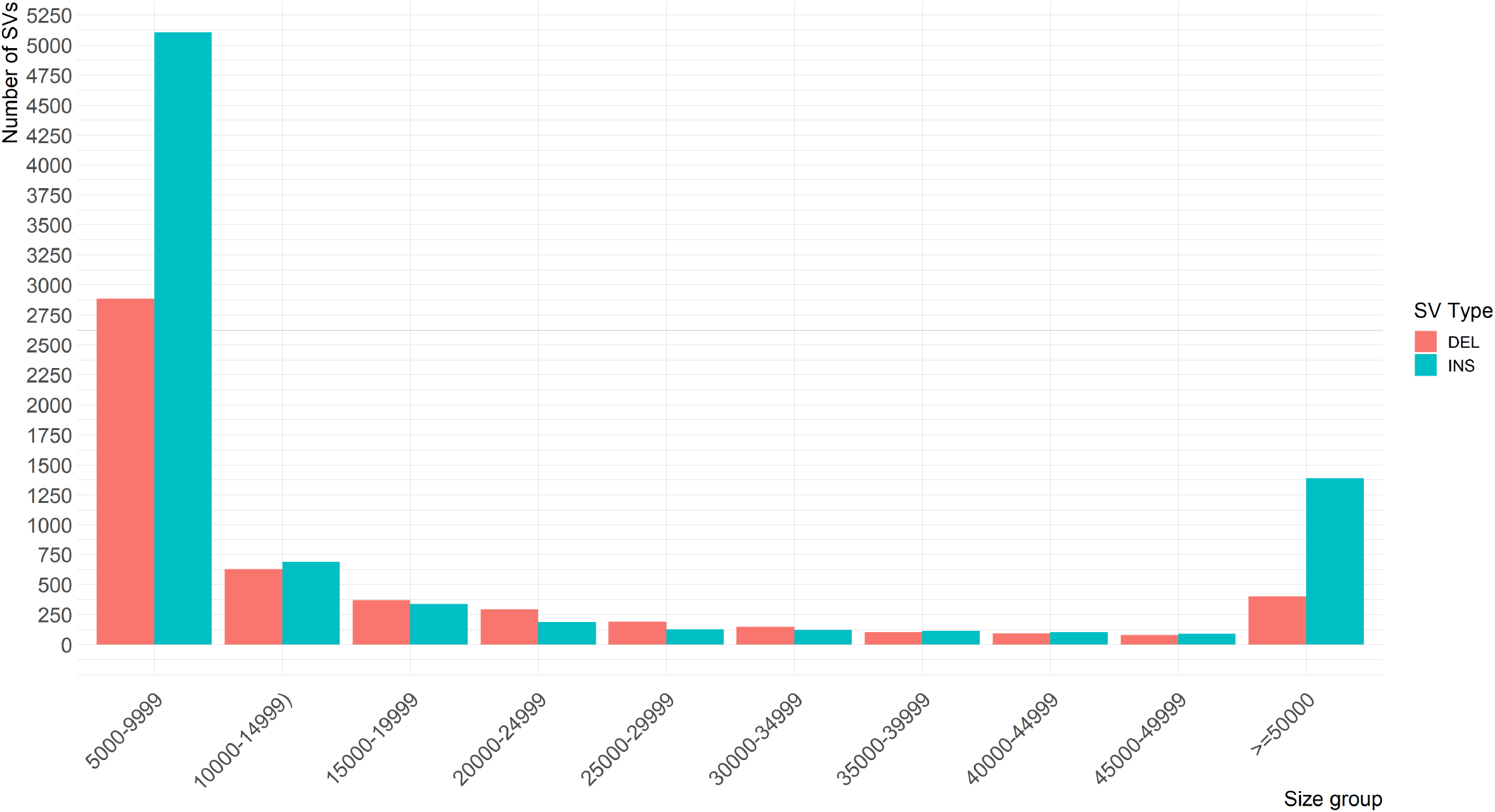
Histogram of the structural variant (SV) sizes. Histogram of the size of the identified SVs in bins of 5Kb.

These SVs longer than 1Kb and of high quality involve a total of 2,656 unique regions, for an estimated total of over 90Mb of non-redundant bases (Supplementary Data 4). This number is comparable to what has been seen for novel sequences (i.e. insertions) using graph genome approaches, where an extra 70Mb and 116Mb of novel sequence were reported on 5 and 4 cattle reference genomes, respectively^28,29^. After merging the filtered variants from all the samples, most of the SVs were found to be private to an individual (Figure 2), consistent with what has been observed in previous studies^1^. Individuals of indicine ancestry (Nelore and Boran) carry almost twice as many SVs relative to the Hereford reference as taurine individuals, further highlighting that the current reference poorly represents these breeds (Figure 2).

**Figure 2:**
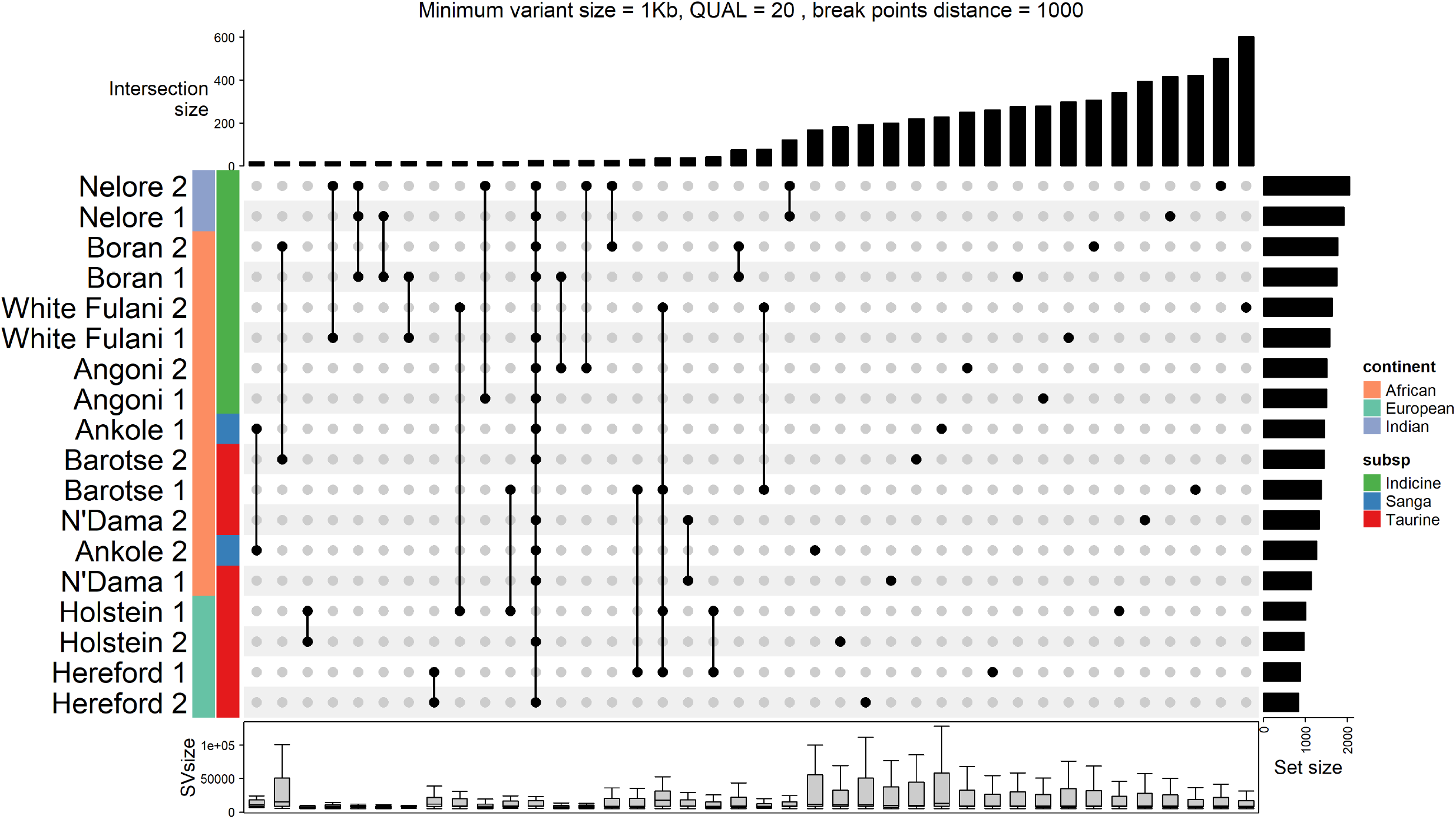
Upset plot of the structural variants. Upset plot of the structural variants by individual for the 40 sets containing the most SVs.

Interestingly, we find that SVs only found in one animal (support = 1; n = 7,445, mean SV length = 85,954.23bp) are generally larger (Wilcoxon rank-sum test P-value = 8.99 * 10^−37^) than the SVs found in more than one animal (support > 1; n = 6,012, mean SV length = 27,747.17bp, Figure 1, Supplementary Figure 1). The list of all SVs, with their position, support and size, are reported in Supplementary Data 5.

Finally, we investigated whether any of the high-quality SVs potentially impact annotated genes. VEP successfully processed 12,999 out of 13,457 SVs (see HTML report on [GitHub](https://htmlpreview.github.io/?https://github.com/evotools/CattleOManalyses/blob/main/Results/VEP/stats_summary.html)). Some variants were too large to be successfully processed by VEP, and other were called as incomplete by VEP. Of these, 6,946 were intergenic, and the remaining 5,934 overlapped5,780 genes and 17,386 transcripts, suggesting the potential for functional variants among the SVs detected. A total of 1,200 SVs putatively overlap a coding sequence. These coding sequences are included in a total of 884 unique gene elements in the cow annotation (Ensembl v105), and of these 483 have an associated gene name (Supplementary Data 6). A total of 292 out of 483 genes had an ID recognized by FUMA^26^. These 292 genes belong to a number of gene sets such as the Hallmark bile acid metabolism and interferon γ and α response sets (Supplementary Figure 2), as well as the olfactory receptor curated gene set (Supplementary Figure 3). All gene set results from FUMA are reported in Supplementary Data 7.

## Usage Notes

Even with the ever-decreasing cost of long read sequencing making it increasingly tractable to call SVs across sets of samples using HTS, validation of the SV calls remain challenging. This compendium of SVs across global cattle breeds provides a validation set called using an independent technology that can be used to assess the quality of cattle SV calls. In fact, optical mapping data has previously been used to validate sequencing based SV calls^28^, and we believe this dataset provides the largest set of optical maps to date for a livestock species.

With many SVs shared across the two animals of each breed, the raw molecules in this dataset can also be used to help scaffold and validate novel assemblies of cattle of breeds closely related to the individuals represented here, potentially reducing the cost of future genome assembly projects.

Unlike most cattle studies, this database is not focused just on European cattle breeds, meaning this will be a valuable resource to researchers across the globe. Importantly, it will allow for SVs to inform the interpretation of results from GWAS and population genetics studies by providing candidate functional variants in relevant regions.

Ultimately, we expect the database to enable further insights into SVs, an understudied class of genetic variation in cattle, giving access to a catalogue of thousands of variants present across multiple breeds worldwide.

## Code availability

The code used in this article were deposited at https://github.com/evotools/CattleOManalyses.

## Acknowledgements

The study was funded by grant BB/R015155/1 to JGDP and Institute Strategic Programme Grant BBS/E/D/10002070 from the Biotechnology and Biological Sciences Research Council (BBSRC).

## Author contribution

J.G.D.P. and T.C. conceived the study, A.T. and J.G.D.P. designed the analyses and A.T. performed them. J.G.D.P. and A.T. prepared the initial manuscript with all authors contributing to subsequent drafts. J.Po., M.C. and T.C. prepared the DNA and Bionano samples. D.W. and P.T. provided samples and expertise for the study. M.E.Z.M., I.M.S., A.F., B.R.F., C.E., E.T.O., D.M., W.A., I.S., J.B.M., G.M. and R.F.K provided samples for the study.

## Notes

https://github.com/evotools/CattleOManalyses/

https://zenodo.org/record/6516933

https://zenodo.org/record/6516993

https://zenodo.org/record/6517172

### Competing Interest Statement

The authors have declared no competing interest.

### Summary of Updates

Changed wrong affiliation for J.B. Muma.

## References

1. Collins, R. L. et al. A structural variation reference for medical and population genetics. Nature 581, 444–451 (2020).

2. Jeffares, D. C. et al. Transient structural variations have strong effects on quantitative traits and reproductive isolation in fission yeast. Nat. Commun. 8, (2017).

3. Liu, G. E. et al. Initial analysis of copy number variations in cattle selected for resistance or susceptibility to intestinal nematodes. Mamm. Genome Off. J. Int. Mamm. Genome Soc. 22, 111–121 (2011).

4. A 12 kb multi-allelic copy number variation encompassing a GC gene enhancer is associated with mastitis resistance in dairy cattle. https://journals.plos.org/plosgenetics/article?id=10.1371/journal.pgen.1009331.

5. Bickhart, D. M. & Liu, G. E. The challenges and importance of structural variation detection in livestock. Front. Genet. 5, (2014).

6. Weischenfeldt, J., Symmons, O., Spitz, F. & Korbel, J. O. Phenotypic impact of genomic structural variation: insights from and for human disease. Nat. Rev. Genet. 14, 125–138 (2013).

7. Chen, L., Chamberlain, A. J., Reich, C. M., Daetwyler, H. D. & Hayes, B. J. Detection and validation of structural variations in bovine whole-genome sequence data. Genet. Sel. Evol. 49, 13 (2017).

8. Couldrey, C. et al. Detection and assessment of copy number variation using PacBio long-read and Illumina sequencing in New Zealand dairy cattle. J. Dairy Sci. 100, 5472–5478 (2017).

9. Bickhart, D. M. et al. Diversity and population-genetic properties of copy number variations and multicopy genes in cattle. DNA Res. 23, 253–262 (2016).

10. Liu, G. E. et al. Analysis of copy number variations among diverse cattle breeds. Genome Res. 20, 693–703 (2010).

11. Mei, C. et al. Copy number variation detection in Chinese indigenous cattle by whole genome sequencing. Genomics 112, 831–836 (2020).

12. Upadhyay, M. et al. Introgression contributes to distribution of structural variations in cattle. Genomics 113, 3092–3102 (2021).

13. Alkan, C., Coe, B. P. & Eichler, E. E. Genome structural variation discovery and genotyping. Nat. Rev. Genet. 12, 363–376 (2011).

14. Pabinger, S. et al. A survey of tools for variant analysis of next-generation genome sequencing data. Brief. Bioinform. 15, 256–278 (2014).

15. Wala, J. A. et al. SvABA: genome-wide detection of structural variants and indels by local assembly. Genome Res. 28, 581–591 (2018).

16. Yuan, Y., Chung, C. Y.-L. & Chan, T.-F. Advances in optical mapping for genomic research. Comput. Struct. Biotechnol. J. 18, 2051–2062 (2020).

17. Lam, E. T. et al. Genome mapping on nanochannel arrays for structural variation analysis and sequence assembly. Nat. Biotechnol. 30, 771–776 (2012).

18. Levy-Sakin, M. et al. Genome maps across 26 human populations reveal population-specific patterns of structural variation. Nat. Commun. 10, (2019).

19. Ellis, J. A. et al. Differentiation antigens on bovine mononuclear phagocytes identified by monoclonal antibodies. Vet. Immunol. Immunopathol. 19, 325–340 (1988).

20. Chan, S. et al. Structural Variation Detection and Analysis Using Bionano Optical Mapping. in Copy Number Variants: Methods and Protocols (ed. Bickhart, D. M.) 193–203 (Springer, 2018). doi:10.1007/978-1-4939-8666-8_16.

21. Danecek, P. et al. Twelve years of SAMtools and BCFtools. GigaScience 10, giab008 (2021).

22. Quinlan, A. R. & Hall, I. M. BEDTools: a flexible suite of utilities for comparing genomic features. Bioinforma. Oxf. Engl. 26, 841–2 (2010).

23. R core team. R: a language and environment for statistical computing. (R Foundation for Statistical Computing, 2021).

24. Gu, Z., Eils, R. & Schlesner, M. Complex heatmaps reveal patterns and correlations in multidimensional genomic data. Bioinformatics 32, 2847–2849 (2016).

25. McLaren, W. et al. The Ensembl Variant Effect Predictor. Genome Biol. 17, 122–122 (2016).

26. Watanabe, K., Taskesen, E., Van Bochoven, A. & Posthuma, D. Functional mapping and annotation of genetic associations with FUMA. Nat. Commun. 8, 1826–1826 (2017).

27. Rosen, B. D. et al. De novo assembly of the cattle reference genome with single-molecule sequencing. GigaScience 9, 1–9 (2020).

28. Talenti, A. et al. A cattle graph genome incorporating global breed diversity. Nat. Commun. 13, 910 (2022).

29. Crysnanto, D., Leonard, A. S., Fang, Z.-H. & Pausch, H. Novel functional sequences uncovered through a bovine multiassembly graph. Proc. Natl. Acad. Sci. 118, e2101056118 (2021).

