## Supplementary figures and images for "Optical mapping compendium of structural variants across global cattle breeds"

### Supplementary Figure 1

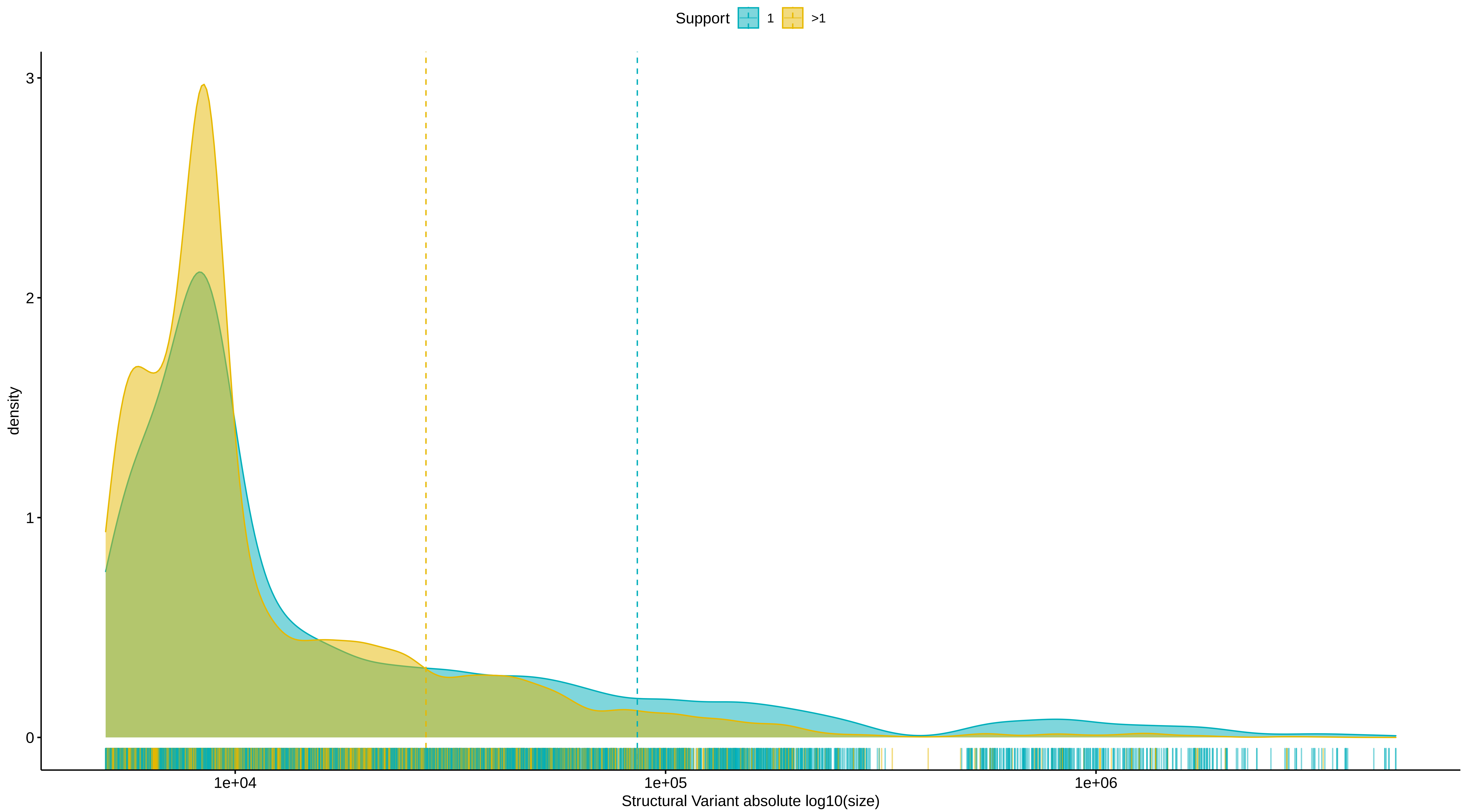

### Supplementary Figure 2

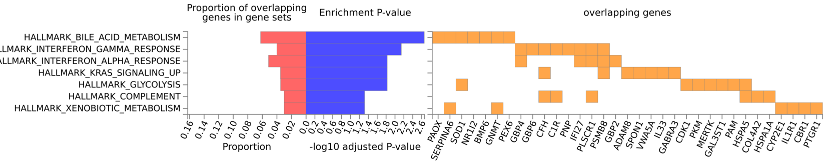

### Supplementary Figure 3

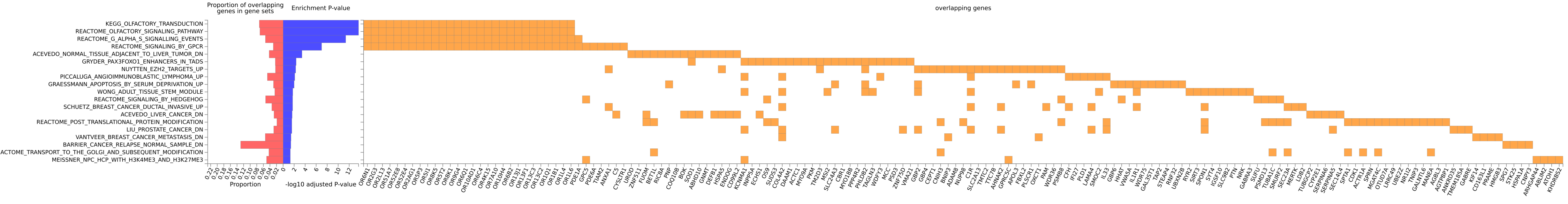
